# Myelin plasticity in ventral tegmental area is required for opioid reward

**DOI:** 10.1101/2022.09.01.506263

**Authors:** Belgin Yalçın, Matthew B. Pomrenze, Karen Malacon, Isabelle J. Chau, Kathryn R. Taylor, Lijun Ni, Daniel Contreras-Esquivel, Robert C. Malenka, Michelle Monje

**Affiliations:** Department of Neurology and Neurological Sciences, Stanford University; Stanford CA 94305; Nancy Pritzker Laboratory, Department of Psychiatry and Behavioral Sciences, Stanford University; Stanford CA 94305; Howard Hughes Medical Institute; Stanford CA 94305

## Abstract

All drugs of abuse induce long-lasting changes in synaptic transmission and neural circuit function that underlie substance use disorders. Here, we demonstrate that dopaminergic neuronal activity-regulated myelin plasticity is a key modulator of dopaminergic circuit function and opioid reward. Oligodendroglial lineage cells respond to dopaminergic neuronal activity evoked by either optogenetic stimulation or by morphine administration specifically within the reward center ventral tegmental area (VTA), but not along the axonal projections in the medial forebrain bundle nor within the target nucleus accumbens (NAc). Genetic blockade of oligodendrogenesis dampens NAc dopamine release dynamics, which is critical for reward learning, and impairs behavioral conditioning to morphine. Our findings identify dopaminergic neuronal activity-regulated myelin plasticity as an important circuit modification that is required for opioid reward.

**One-Sentence Summary:** Activity-dependent myelin plasticity in the ventral tegmental area modulates dopaminergic circuit function and opioid reward

## Main Text

Motivated behavior - critical for adapting to the environment and therefore for animal survival - depends on proper function of reward circuitry. Drugs of abuse induce maladaptive and persistent modifications in reward circuitry, facilitating the development of addictive behaviors. All drugs of abuse, including opioids, target the midbrain dopaminergic (DA) reward system and induce lasting changes in synaptic transmission and neural circuit function (*1–5*). Morphine, a natural opioid found in opium, triggers synaptic plasticity and alters neuronal function in the ventral tegmental area (VTA) and nucleus accumbens (NAc), two key structures of the DA reward system, promoting a pathological form of reward learning (*1–5*). These experience-dependent alterations of reward system function are critical for shaping drug-induced behavioral changes and hence the development of substance use disorders (*1–5*).

While a significant role for microglia and astrocytes in these neural circuit modifications is becoming increasingly apparent (*6–10*), contributions of oligodendroglial lineage cells and myelin to reward circuit adaptation in health and maladaptation in addictive states remain unknown. However, oligodendroglial lineage cells are a glial cell type particularly well positioned to contribute to structural and functional changes in addiction-related neural circuits. Oligodendrocytes generate myelin that ensheaths axons to modulate conduction velocity (*11, 12*) and provide metabolic support to axons (*13, 14*), therefore playing a fundamental role in shaping neural transmission. Neuronal activity can regulate myelination during development (*15, 16*) and myelin plasticity in adulthood through OPC proliferation, oligodendrogenesis and de novo myelination by new oligodendrocytes and through myelin remodeling by existing oligodendrocytes (*17–22*). Even small changes in myelination can tune circuit dynamics (*23–25*) and consequently influence cognition and behavior (*17, 26–31*). These circuit-specific, activity-regulated myelin changes appear to occur only in distinct neuronal types (*17, 22*). Thus, which neuronal subtypes, in which circuits and with which patterns of activity elicit oligodendroglial responses remain to be fully understood. Furthermore, much remains to be learned about how activity-regulated myelination may become maladaptive and contribute to the pathophysiology of neurological (*32*) or psychiatric (*33*) diseases.

Here, we explore the role of myelin plasticity in morphine-elicited reward and morphine-elicited DA release dynamics in the NAc. Our results identify myelin plasticity as a previously unappreciated feature of the activity-dependent modifications of reward circuit function that critically contribute to the behavioral reinforcing effects of opioids.

## Results

### DA neuron activity-regulated oligodendrogenesis and myelination

We leveraged *in vivo* optogenetic strategies to examine whether DA neuron activity regulates oligodendroglial cell dynamics in reward circuitry. To directly stimulate DA neurons, we injected adeno-associated virus carrying Cre-inducible channelrhodopsin2 (AAV-Ef1a-DIO-ChR2::YFP) or YFP (AAV-Ef1a-DIO-YFP) into the VTA of Dat-Cre mice (**Fig. 1A; Sup Fig. 1A**). This method achieves DA neuron-specific expression of ChR2 or YFP in VTA, where DA neuron cell bodies are localized, and also in their axonal projections toward other brain regions including the major downstream target, nucleus accumbens (NAc). Localization of ChR and YFP to DA neurons were confirmed by tyrosine hydroxylase (TH) co-localization in VTA (**Sup Fig. 1B**). Optogenetic stimulation of VTA at 30 Hz increased cfos signal co-localizing with ChR^+^ dopaminergic neurons after 90 minutes, indicating successful optogenetic targeting (**Sup Fig. 1C, D, and E**).

**Fig. 1.**
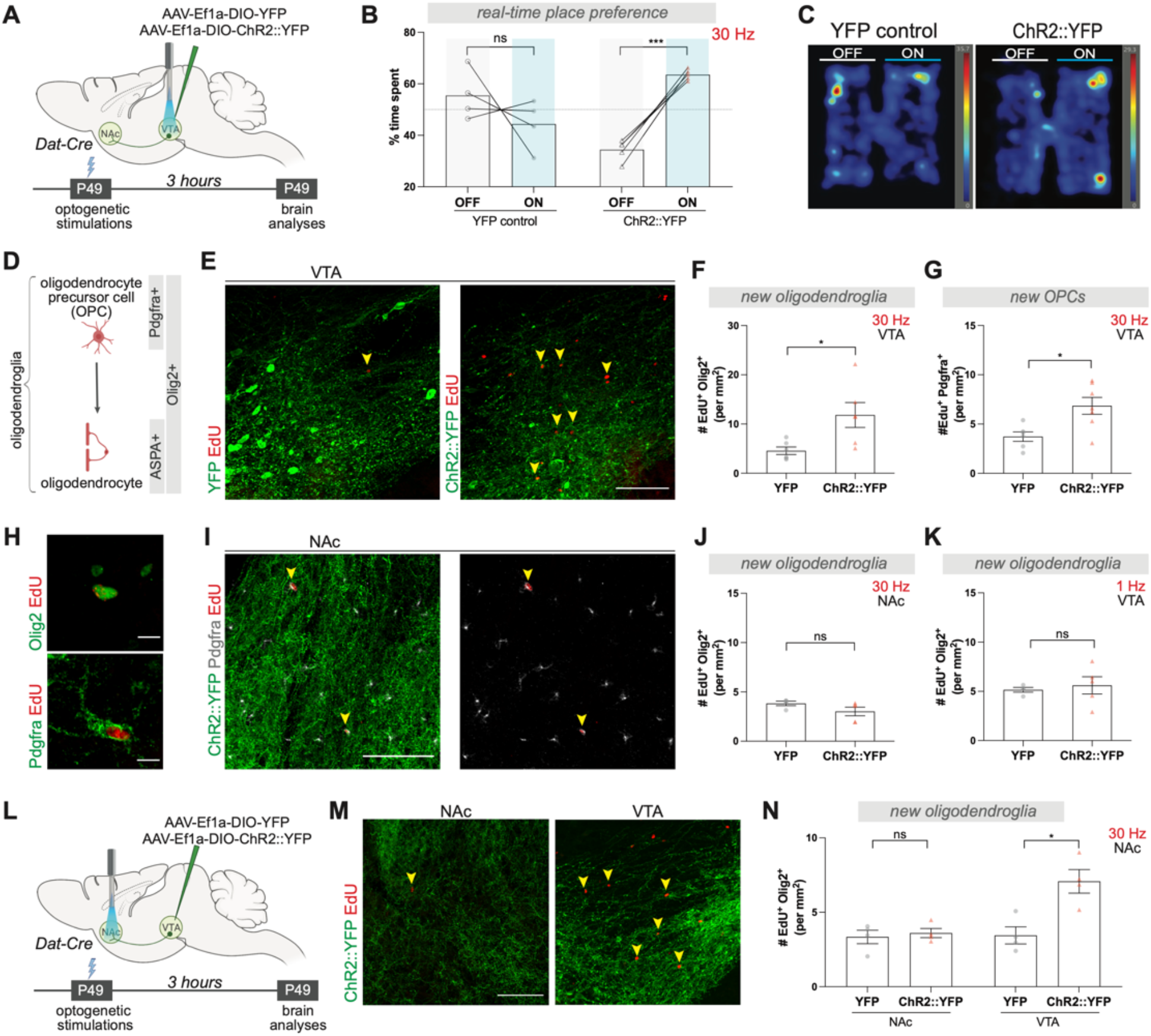
Reward circuitry DA neuron activity increases OPC proliferation specifically in VTA. **(A)** Schematic of experimental paradigm for acute DA neuron stimulation in VTA; 30 minutes optogenetic stimulation followed by brain analysis after 3 hours. **(B)** Real-time place preference test at 30Hz. Percentage of time spent in each chamber compartment paired with and without laser stimulation for YFP control and ChR2::YFP mice. 30 hz optogenetic stimulation of DA neurons increases place preference (YFP, n=4 mice; ChR2::YFP, n=5 mice) **(C)** Representative heat maps illustrating the time spent in both compartments of the chamber during the real-time place preference test. **(D)** Schematic of oligodendroglial lineage and specific markers; Olig2 is an oligodendroglial lineage marker, Pdgfra marks OPCs, and ASPA marks mature oligodendrocytes. **(E)** Confocal micrographs of proliferating (EdU^+^) cells in VTA, and **(I)** in NAc, scale bars= 100 μm **(F)** 30 hz VTA stimulation of DA neurons increases new oligodendroglia density (Olig2^+^ EdU^+^), and **(G)** new OPC density (Pdgfra^+^ EdU^+^) in VTA (YFP, and ChR2::YFP, n=6 mice). **(H)** Confocal micrographs show a new oligodendroglial cell (Olig2^+^ EdU^+^), and an OPC (Pdgfra^+^ EdU^+^), scale bars= 10 μm. **(J)** New oligodendroglia density (Olig2^+^ EdU^+^) in NAc does not change after 30 hz VTA stimulation or **(K)** in VTA after 1 hz VTA stimulation (YFP, n=4 mice; ChR2::YFP, n=5 mice). **(L)** Schematic of experimental paradigm for acute DA neuron stimulation in NAc; 30 minutes optogenetic stimulation followed by tissue analysis after 3 hours. **(M)** Confocal micrographs showing proliferating cells (EdU^+^) in NAc and VTA after 30 hz NAc stimulation near DA neurons ChR2::YFP (green), arrowheads denote new cells, scale bars= 100 μm. **(N)** 30 hz NAc stimulation of DA neurons increases new oligodendroglia density (Olig2^+^ EdU^+^) in VTA but not in NAc (YFP, and ChR2::YFP, n=4 mice), scale bars= 100 μm. In all graphs data points represent each mouse. **B**, paired two-tailed t-test for each condition, **H, I, J, K, N,** unpaired two-tailed t-test, ns (not significant) p>0.5, *p<0.05, **p<0.01, ***p<0.001. Data shown as mean, error bars indicate SEM.

DA neurons exhibit different activity modes; “phasic firing” indicates synchronized bursts of action potentials at frequencies above 10 Hz, whereas “tonic firing” refers to spontaneous activity between 0.2-10 Hz (*34, 35*). Because tonic and phasic firing are associated with different levels of DA release and can mediate different aspects of motivated behavior, we tested both of these activity modes. With stimulation of VTA at 30 Hz, ChR2-expressing mice showed a strong preference for the stimulation-paired chamber in a real time place preference assay, confirming a rewarding effect of phasic DA neuron activation (*35*) (**Fig. 1B, and C**). To assess whether this phasic activity affects oligodendroglial lineage cells (**Fig. 1A**), we used cell identity markers; specifically, all oligodendroglial cells express Olig2, while Pdgfra marks only oligodendroglial precursor cells (OPCs), and mature oligodendrocytes are labelled by ASPA (**Fig. 1D**). In addition, to mark dividing cells, we administered the thymidine analog 5-ethynyl-2’-deoxyuridine (EdU) at the beginning of each optogenetic stimulation session. Acute phasic optogenetic stimulation of VTA DA neurons at 30 Hz intermittently over 30 minutes (eight pulses of 5 ms light flashes delivered every 5 seconds) significantly increased proliferating oligodendroglial lineage cell (EdU^+^ Olig2^+^) density and promoted proliferation of OPCs (EdU^+^ Pdgfra^+^) in VTA within three hours (**Fig. 1E, F, G, and H**). Notably, there was no increase in new oligodendroglia (EdU^+^ Olig2^+^) in the NAc, where axons and presynaptic terminals of DA neurons are located (**Fig. 1I and J**). By contrast, tonic stimulation, which does not cause real time reward or aversion (*35*), at 1 Hz for 30 minutes (25 pulses of 15 ms light flashes delivered every 1 minute) did not change the proliferation of oligodendroglial cells (EdU^+^ Olig2^+^) in VTA, suggesting that the firing mode of DA neurons is critical for the activity-regulated oligodendroglial response (**Fig. 1K**).

As we observed a proliferative increase only in VTA OPCs (**Fig. 1G**), we next investigated whether the oligodendroglial response to DA neuron activity is brain region-specific. We once again infected VTA DA neurons with ChR2 but placed the fiber optic over the NAc and stimulated at 30 Hz for 30 minutes as above (**Fig. 1L**). Phasic stimulation of VTA DA axon terminals in the NAc failed to increase proliferating oligodendroglial lineage cells (EdU^+^ Olig2^+^) locally in NAc. However, we again observed oligodendroglial cell proliferation in the VTA (**Fig. 1M and N**), presumably because of antidromic activation of VTA DA neuron cell bodies (*36*). Taken together, these data demonstrate that phasic VTA DA neuron activity promotes OPC proliferation specifically in VTA.

Next, we evaluated the cell fate of the new oligodendroglial cells proliferating in response to phasic DA neuron activity. To achieve a more sustained increase in neuronal activity, we stimulated DA neurons at 30 Hz for 10 minutes a day for 7 days and analyzed the brains 4 weeks after optogenetic stimulations (**Fig. 2A**). We examined new oligodendroglial lineage cells across the VTA→NAc pathway, including in the NAc and along medial forebrain bundle (MFB), through which DA axons project. Neither NAc (**Fig. 2B**) nor MFB (**Fig. 2C**) showed a change in new oligodendroglia (EdU^+^ Olig2^+^) after a prolonged increase in DA neuron activity. However, there was a substantial increase in new oligodendrocyte (EdU^+^ ASPA^+^) density in the VTA (**Fig. 2D and E**), once again highlighting the specificity for the activity-regulated oligodendroglial response in the VTA. This specificity may reflect the regional heterogeneity of OPCs across the brain (*37*), and/or the microenvironmental differences - such as proximity to DA cell bodies versus axons (*38*), or may reflect neuron subtype-specific biology (*17, 22*).

**Fig. 2.**
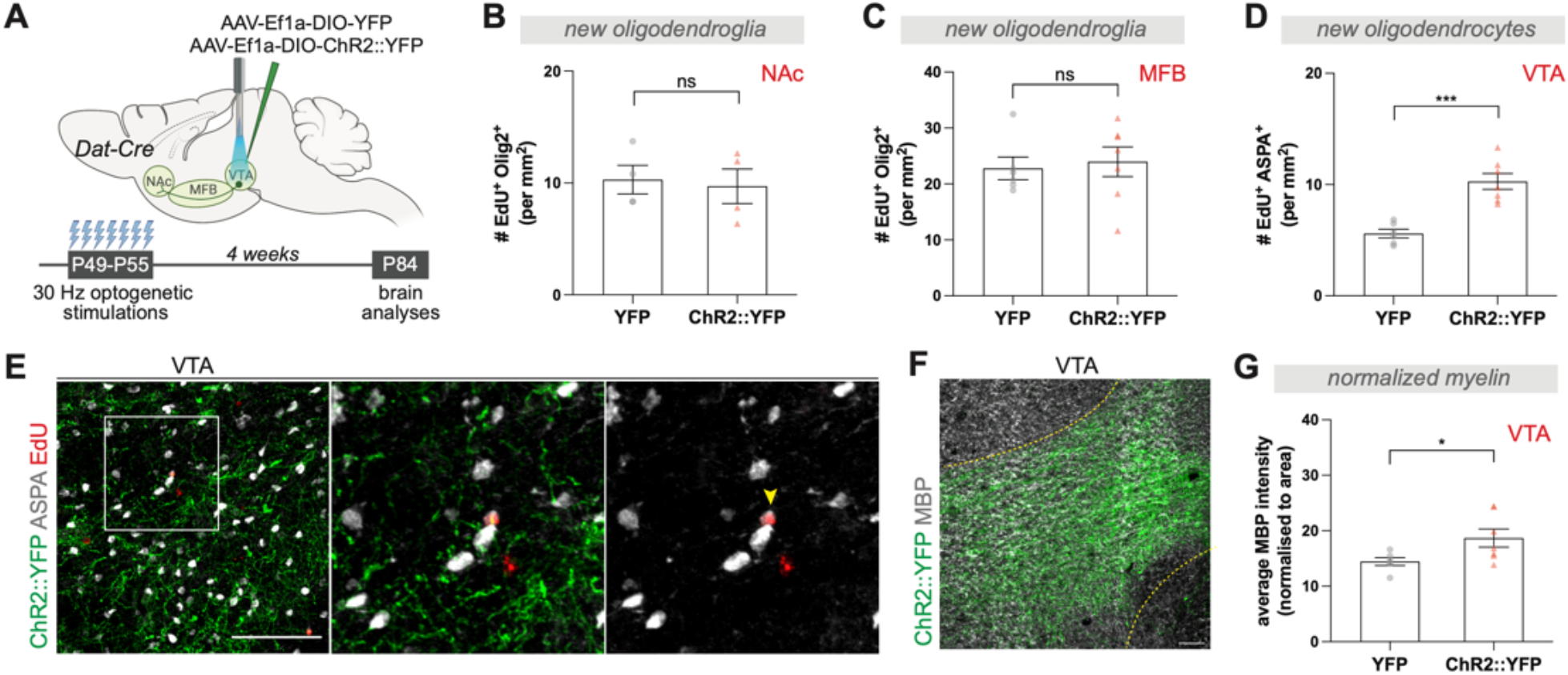
DA neuron activity regulates oligodendrogenesis in VTA. **(A)** Schematic of experimental paradigm for chronic DA neuron stimulation in VTA; 30 hz optogenetic stimulation for 10 minutes a day for 7 days, followed by brain analysis 4 weeks after this paradigm. **(B)** Chronic 30 hz DA neuron stimulation do not affect new oligodendroglia density (Olig2^+^ EdU^+^) in NAc (YFP, n=4 mice; ChR2::YFP, n=4 mice) or **(C)** MFB but **(D)** increases new oligodendrocyte density (ASPA^+^ EdU^+^) in VTA (YFP, n=6 mice; ChR2::YFP, n=7 mice). **(E)** Confocal micrographs showing new oligodendrocytes (ASPA^+^ EdU^+^) in VTA after chronic stimulation, arrowhead denotes a new oligodendrocyte with overlapping signals of EdU (red) and ASPA (gray), scale bar= 50 μm. **(F)** Confocal micrograph showing myelination in VTA, yellow dotted line marks VTA border, DA neurons expressing ChR2::YFP (green) and MBP (gray), scale bar= 50 μm **(G)** Chronic 30 hz DA neuron stimulation increases normalized myelin (ratio of average MBP intensity over the area of DA neurons in VTA) (YFP, n=6 mice; ChR2::YFP, n=7 mice). Unpaired two-tailed t-test, ns (not significant) p>0.5, *p<0.05, ***p<0.001. Data shown as mean, error bars indicate SEM.

As a mature oligodendrocyte can generate between 20-50 myelin sheaths, even small changes in oligodendrocyte numbers can exert significant effects on myelination and conduction velocity (*25, 39*). While electron microscopy analyses are challenging in this anatomical region, to assess potential myelination changes in VTA as a result of activity-regulated oligodendrogenesis, we measured myelin basic protein (MBP) over ChR2^+^ and YFP^+^ areas. After a prolonged phasic DA neuron stimulation, MBP intensity was significantly increased in the VTA of ChR2 animals compared with those expressing YFP (**Fig. 2F and G**), implying a change in myelination within VTA, potentially involving either the proximal axonal segments of DA neurons or local interneurons in VTA.

### Morphine promotes oligodendrogenesis in VTA

Drugs of abuse initiate persistent adaptations in reward circuitry, laying foundations for drug-seeking behaviors and increased sensitivity to drug cues. We investigated whether acute morphine administration alters oligodendrogenesis as a part of these initial neural circuit adaptations. Wildtype mice were injected systemically with one dose of morphine (10 mg/kg, IP) or saline vehicle as a control, and EdU to mark the dividing cells (**Fig. 3A**). Three hours later, the VTA exhibited an increase in the number of new OPCs (EdU^+^ Pdgfra^+^) compared to saline-treated controls (**Fig. 3B and C**). Similar to the oligodendroglial response to acute phasic optogenetic stimulation of DA neurons (**Fig. 1G and N**), this OPC increase was specific to the VTA (**Fig. 3C**). Oligodendroglial cells express k-opioid receptors (*40, 41*), therefore we tested whether morphine directly affects OPC dynamics *in vitro.* Incubation of OPCs with different concentrations of morphine did not change OPC proliferation (EdU^+^ Pdgfra^+^), in comparison to proliferative media positive control conditions, suggesting that the effects on oligodendroglial cells in the brain microenvironment is not a direct effect of the drug, but rather mediated through cell-cell interactions (**Sup Fig. 2A and B**). We also did not detect a change in OPC proliferation (EdU^+^ Pdgfra^+^) in response to varying DA concentrations *in vitro*, indicating that oligodendroglial cells are not simply responding to neurotransmitter release in the VTA, a finding consistent with the lack of OPC response in NAc where DA neurons release DA. These data suggest that the effect of morphine on oligodendroglial proliferation is chiefly via indirect mechanisms such as neuron-OPC interactions; as morphine increases DA neuron activity, this may in-turn affect oligodendroglial cells.

**Fig. 3.**
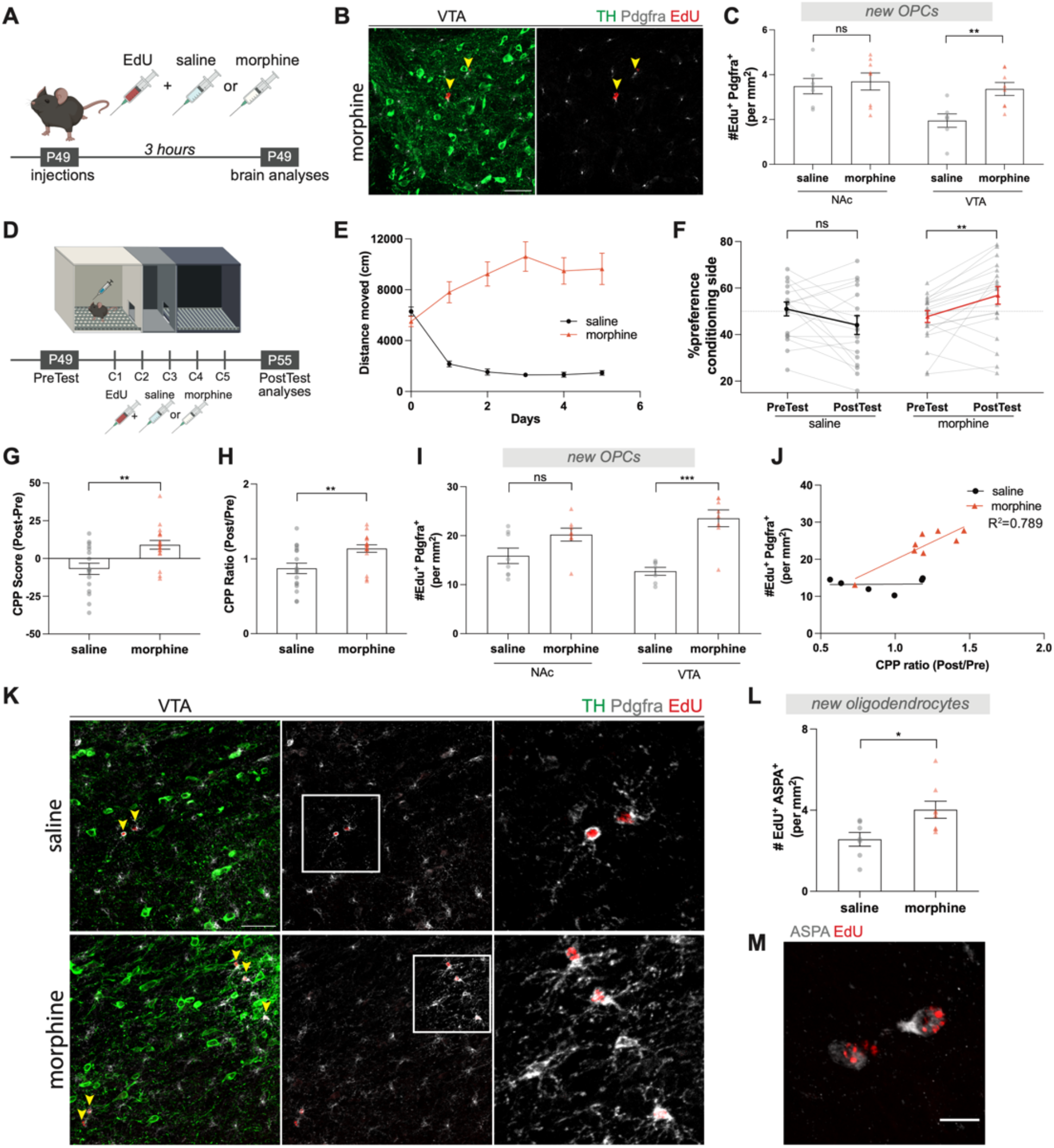
Morphine promotes oligodendrogenesis in VTA. **(A)** Schematic of experimental paradigm, mice groups injected with saline and EdU, or morphine (10mg/kg) and EdU at 7 weeks, and brains analyzed after 3 hours. **(B)** Confocal micrographs showing proliferating OPCs in VTA 3 hours after single dose morphine injection, arrowheads denote new OPCs (Pdgfra^+^ EdU^+^), TH marks DA neurons (green), Pdgfra^+^ (gray), EdU^+^ (red). **(C)** Single dose of morphine increases new OPC density (Pdgfra^+^ EdU^+^) in VTA but not in NAc (saline, n=7 mice; morphine, n=8 mice). **(D)** Schematic of experimental paradigm for conditioned place preference test (CPP), mice groups injected with saline and EdU, or morphine (10mg/kg) and EdU during conditioning sessions, and brains analyzed after PostTest. **(E)** Morphine conditioning increases locomotor sensitivity (saline, n=6 mice; morphine, n=12 mice) over conditioning days, and **(F)** induces a robust a place preference for the conditioning chamber. Graph shows the percentage of time spent in conditioning chamber, comparing PreTest and PostTest (saline, n=17 mice; morphine, n=19 mice). **(G)** Morphine conditioning increases CPP Score (PostTest preference – PreTest preference) and **(H)** CPP ratio, (ratio of PostTest preference to PreTest preference for conditioning chamber) (saline, n=17 mice; morphine, n=18 mice). **(I)** Morphine conditioning increases new OPC density (Pdgfra^+^ EdU^+^) in VTA but not in NAc (saline, and morphine, n=8 mice). **(J)** Number of new OPCs strongly correlates (R^2^=0.789) with CPP ratio (saline, n=7 mice; morphine, n=10 mice). **(K)** Confocal micrographs showing proliferating OPCs in VTA after morphine CPP, arrowheads denote new OPCs (Pdgfra^+^ EdU^+^), TH marks DA neurons (green), Pdgfra^+^ (gray), EdU^+^ (red), scale bar= 50 μm. **(L)** Morphine conditioning increases new oligodendrocytes (ASPA^+^ EdU^+^) in VTA (saline, n=7 mice; morphine, n=8 mice). **(M)** Confocal micrograph showing a new oligodendrocyte after morphine exposure with overlapping signals of EdU (red) and ASPA (gray), scale bar= 10 μm. **F,** paired t-test, **J**, simple linear regression, **C, G, H, I, L,** unpaired two-tailed t-test, ns (not significant) p>0.5, *p<0.05, ***p<0.001. Data shown as mean, error bars indicate SEM.

Opioids recruit different components of reward circuits to adapt and mediate reward learning behavior. We next investigated oligodendroglial cells after morphine-induced conditional place preference (CPP), a form of Pavlovian conditioning in which a neutral context or cue becomes associated with the rewarding or aversive properties of a stimulus (*42*) (**Fig. 3D**). Wildtype mice were conditioned with morphine or saline for 5 days and administered EdU during conditioning sessions to concurrently trace new OPCs. As expected, morphine-treated mice exhibited increasing locomotor activity in response to repeated injections, a form of behavioral sensitization (*43*) (**Fig. 3E**). They also exhibited a preference for the morphine-paired conditioning chamber, that is conditioned place preference (CPP) (**Fig. 3F, G, and H**). These behavioral changes were accompanied by a substantial increase in the number of new VTA OPCs (EdU^+^ Pdgfra^+^), but no increases in NAc OPCs (**Fig. 3I and K**). We found a significant correlation between number of new OPCs (EdU^+^ Pdgfra^+^) in VTA and the CPP ratio, suggesting that morphine may recruit elements of experience-dependent myelination during this reward learning task (**Fig. 3J**). Concordantly, differentiation of these newly proliferated, EdU-marked OPCs in VTA into new oligodendrocytes (EdU^+^ ASPA^+^) was already evident by the end of this experimental paradigm (**Fig. 3L and M**). Together, these data demonstrate that both acute and chronic administration of morphine promotes oligodendrogenesis exclusively in VTA. Furthermore, this VTA-specific oligodendroglial response and the findings that morphine or DA alone are not sufficient to produce this response suggests that morphine-evoked oligodendrogenesis in VTA is an indirect result of enhanced reward circuit activity.

### Loss of oligodendrogenesis blocks morphine reward learning

Neuronal activity-regulated myelination is required for certain forms of learning in healthy brain (*28, 30, 31*). Because morphine-induced CPP behavior is a form of associative reward learning, we hypothesized that adaptive oligodendrogenesis in the VTA is necessary for forming associations between the morphine administrations and the conditioning chamber context. To conditionally block oligodendrogenesis, we deleted myelin regulatory factor (Myrf), a transcriptional factor essential for oligodendroglial differentiation, in oligodendrocyte precursor cells by crossing inducible Pdgfra-Cre^ERT^ mice to floxed *Myrf* (*Myrf*^fl/fl^) mice generating *Myrf^-/-^; Pdgfra-Cre^ERT^* (referred as *Myrf^-/-^* hereafter) (*44*). Deletion of Myrf is achieved upon tamoxifen administration, and therefore these precursors cannot differentiate into oligodendrocytes and instead undergo apoptosis, yet pre-existing oligodendrocytes and myelin remain unaffected (*28*). After 15 weeks of Myrf loss, we analyzed new oligodendroglial lineage cell generation in the myelin-rich corpus callosum to determine the baseline effects of Myrf deletion (**Sup Fig. 3A**). Within 4 weeks after EdU labeling, a significant reduction in new oligodendroglial lineage cell (EdU^+^ Olig2^+^) density was apparent (**Sup Fig. 3B**). Therefore, tamoxifen administration at 7-weeks of age, immediately prior to the CPP assay, prevented further generation of oligodendrocytes in *Myrf^-/-^* animals during morphine conditioning, but this differentiation process central to oligodendrogenesis remained intact in the control Cre^-^ littermates (**Fig. 4A**).

**Fig. 4.**
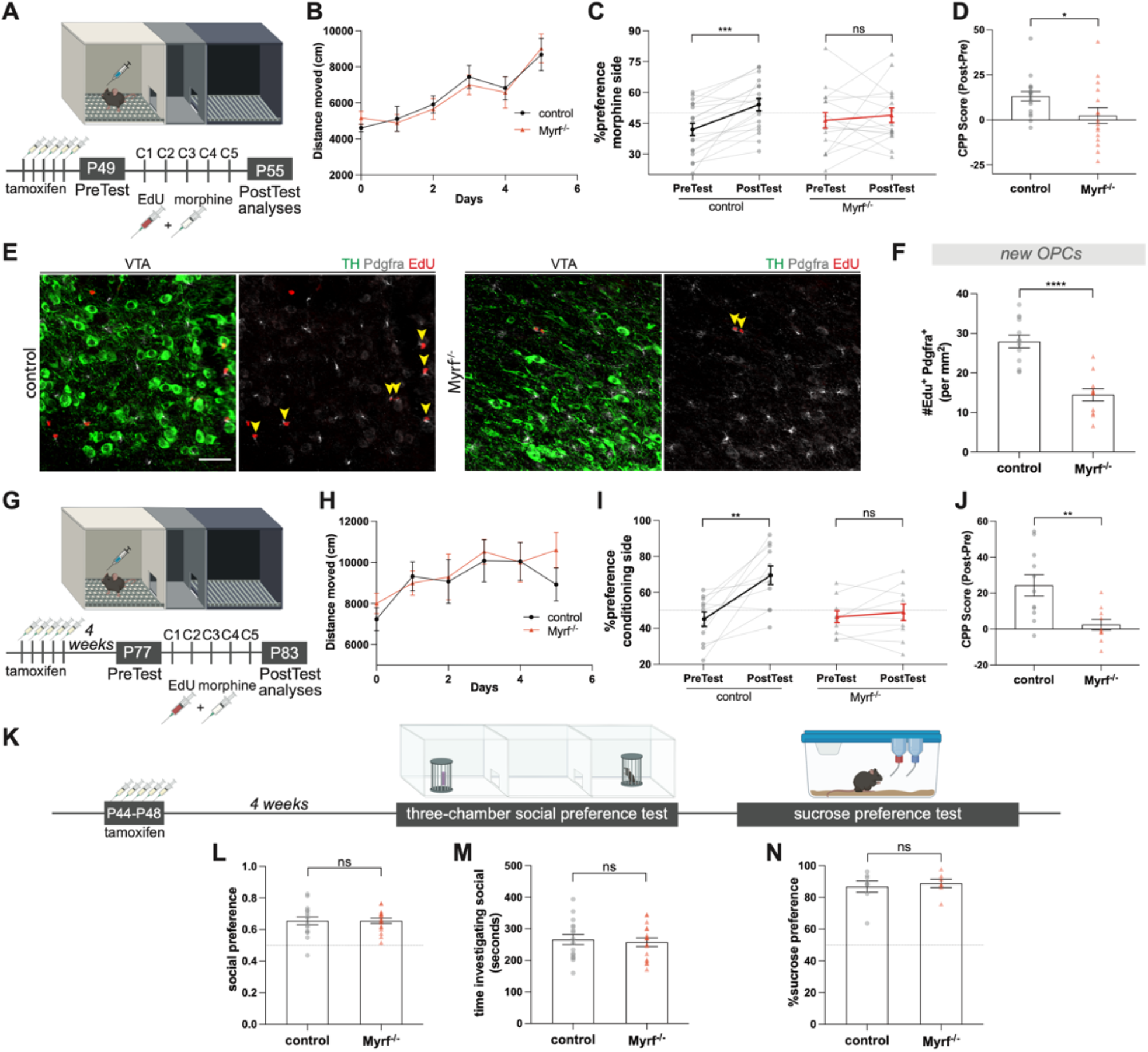
Conditional blockade of oligodendrogenesis abrogates morphine-induced reward learning. **(A)** Schematic of experimental paradigm for *Myrf^-/-^* (*Myrf^-/-^; Pdgfra-Cre^ERT^*) and littermate controls; conditional knock-out performed with tamoxifen administration at 7-weeks of age, immediately prior to CPP, and **(G)** 4-weeks before CPP. Mice groups injected with morphine (10mg/kg) and EdU during conditioning sessions, and brains analyzed immediately after PostTest. **(B)** and **(H)** Both control and *Myrf^-/-^* mice exhibits increased locomotor sensitivity during morphine conditioning (**B**, control, n=20 mice; *Myrf^-/-^*, n=16 mice; **H**, control, n=11 mice; *Myrf^-/-^*, n=10 mice) **(C)** and **(I)** Control mice acquire a place preference for the morphine conditioning chamber, whereas *Myrf^-/-^* mice do not show a strong preference. Graph shows the percentage of time spent in conditioning chamber, comparing PreTest and PostTest (**C**, control, n=19 mice; *Myrf^-/-^*, n=16 mice; **I**, control, n=11 mice; *Myrf^-/-^*, n=10 mice). **(D)** and **(J)** *Myrf^-/-^* animals show decreased CPP Score (PostTest preference – PreTest preference) compared to controls **(E)** Confocal micrographs showing proliferating OPCs in VTA after morphine CPP, arrowheads denote new OPCs (Pdgfra^+^ EdU^+^), TH marks DA neurons (green), Pdgfra^+^ (gray), EdU^+^ (red), scale bar= 50 μm. **(F)** Loss of Myrf decreases number of new OPCs in VTA after morphine CPP (control, n=13 mice; *Myrf^-/-^*, n=11 mice). **(K)** Schematic of experimental paradigm for *Myrf^-/-^* (*Myrf*^-/-^; *Pdgfra-Cre^ERT^*) and littermate controls; conditional knock-out performed with tamoxifen administration at 7-weeks of age, 4-weeks prior to behavioral tests. **(L)** Both control and *Myrf^-/-^* mice show strong preference for the social subject during a three-chamber social preference test and **(M)** spent similar time investigating social subject (control, n=16 mice; *Myrf^-/-^*, n=17 mice). **(N)** Both control and *Myrf^-/-^* mice show strong preference for sucrose over water (control, n=8 mice; *Myrf^-/-^*, n=7 mice). **C, I** paired two-tailed t-test, **D, F, J, L, M, N** unpaired two-tailed t-test, ns (not significant) p>0.5, *p<0.05, **p<0.01, ***p<0.001, ****p<0.0001. Data shown as mean, error bars indicate SEM.

In the CPP experiments, *Myrf^-/-^* animals displayed increased locomotor sensitization similar to the controls, yet morphine CPP was significantly abrogated (**Fig. 4B-D**). This finding suggests that oligodendrogenesis is necessary for morphine reward learning, but not for locomotor sensitization. Consistent with this conclusion, *Myrf^-/-^* animals showed a substantially reduced oligodendroglial response (EdU^+^ Pdgfra^+^) in the VTA compared with littermate controls (**Fig. 4E and F**).

Because Myrf loss and consequent abrogation of oligodendrogenesis is linked to impairments in hippocampal memory consolidation over time (at 4 weeks, but not at 1 day, after Myrf recombination) (*30, 31*), we tested *Myrf^-/-^* animals to determine whether the observed abrogation of CPP may be due to general memory deficits by performing a memory task within the timeline that we perform morphine CPP experiments. One week following tamoxifen-induced Myrf deletion, coinciding with the timeframe of the CPP paradigm, we tested mice in the novel object recognition task (NORT), where mice are introduced with two identical objects and then exposed to one familiar and one novel object 24 hours later (*45*) (**Sup Fig. 3C**). Healthy animals recognize the old object and spend more time exploring the novel object, while mice with memory deficits cannot discriminate between the familiar and novel objects and spend similar time exploring each object (*45*). At 24 hours, both control and *Myrf^-/-^* animals explored the novel object for more time than the familiar object (**Sup Fig. 3D**). These results demonstrate that, at this time point examined – one week after Myrf recombination – loss of oligodendrogenesis does not cause significant general memory deficits.

We also tested the prolonged effects of blocking oligodendrogenesis by assessing morphine CPP 4 weeks after tamoxifen-induced Myrf deletion (**Fig 4G**). Replicating the data at the one-week timepoint, both control and *Myrf^-/-^* animals displayed locomotor sensitization to morphine (**Fig. 4H**). Control animals acquired a robust preference for the morphine-paired conditioning chamber whereas for *Myrf^-/-^* animals this response was significantly abrogated (**Fig. 4I and J**).

To address whether blocking oligodendrogenesis can alter behaviors in response to natural rewards, we tested *Myrf^-/-^* animals 4 weeks after deletion for sociability and sucrose preference (**Fig. 4K**). In the three-chamber social preference test, we found that both control and *Myrf^-/-^* animals exhibited comparable preferences towards a same-sex juvenile conspecific over an unanimated object (**Fig. 4L**), reflected in similar amounts of time investigating the juvenile (**Fig. 4M**). These data demonstrate that blocking oligodendrogenesis at this timepoint and age does not affect general sociability. We also tested preference for a natural reward, sucrose, to determine whether the abrogation of a preference for morphine in *Myrf^-/-^* animals could be due to anhedonia-like effects (*46*). Both control and *Myrf^-/-^* animals showed a strong preference towards sucrose solution over water, indicating that blocking oligodendrogenesis did not cause a general decrease in reward processing (**Fig. 4N**). Taken together these data demonstrate that oligodendrogenesis is required for morphine-induced reward learning, but not for intrinsic reward behaviors.

### Maladaptive oligodendrogenesis alters DA release dynamics in NAc

Myelin adapts with changing neuronal activity and mediates neural network coordination to ensure proper spike-time arrival for optimal network function (*23*). We hypothesized that morphine-induced oligodendrogenesis synchronizes and strengthens DA neuron activity, thereby optimizing DA release dynamics in projection targets to facilitate reward learning and memory retrieval. To test this possibility, we expressed the genetically encoded fluorescent DA sensor GRAB_DA_ (*47*) in the NAc medial shell and recorded real-time endogenous DA dynamics in mice undergoing the morphine CPP procedure. We performed recordings during the PreTest and PostTest to determine whether DA release dynamics adapt with morphine reward learning (**Fig. 5A, and B**). In line with our previous findings (**Fig. 4**), both control and *Myrf^-/-^* animals displayed locomotor sensitization over the morphine conditioning days, and control animals acquired a strong preference to the morphine-paired conditioning chamber which was absent in *Myrf^-/-^* animals (**Fig. 5C and D**). During the PreTest baseline recording prior to reward conditioning, both control and *Myrf^-/-^* animals showed small spontaneous NAc DA transients during exploration of the chamber to be paired with morphine, likely due to novelty (**Fig. 5E and F**). After 5 days of conditioning with morphine, control animals displayed a significant increase in NAc DA levels upon their first entry into the morphine-paired chamber during the PostTest, an expression of reward expectation for the previous morphine experience associated with the chamber (*48*) (**Fig. 5E**). By contrast, *Myrf^-/-^* animals showed a significantly blunted DA response upon entry, similar to their PreTest levels, suggesting an absence of an association between the chamber and the previous morphine rewards (**Fig. 5E, G, and H**). During the PreTest, DA release in *Myrf^-/-^* animals was similar to controls, illustrating that *Myrf^-/-^* animals do not exhibit lower baseline levels of DA (**Fig. 5F**). After morphine conditioning, control animals associate the conditioning chamber with the morphine reward, and exhibit a significant increase in NAc DA levels when they enter into the conditioning chamber with a reward expectation during the PostTest (**Fig. 5G**). Strikingly, this learned DA response is lacking in oligodendrogenesis-deficient *Myrf^-/-^* animals, reflected in the difference score of fluorescence from the PostTest and PreTest (**Fig. 5G, and H**). Overall, these results suggest that morphine-induced oligodendrogenesis in the VTA promotes DA release dynamics in the NAc, critical for the reward learning.

**Fig. 5.**
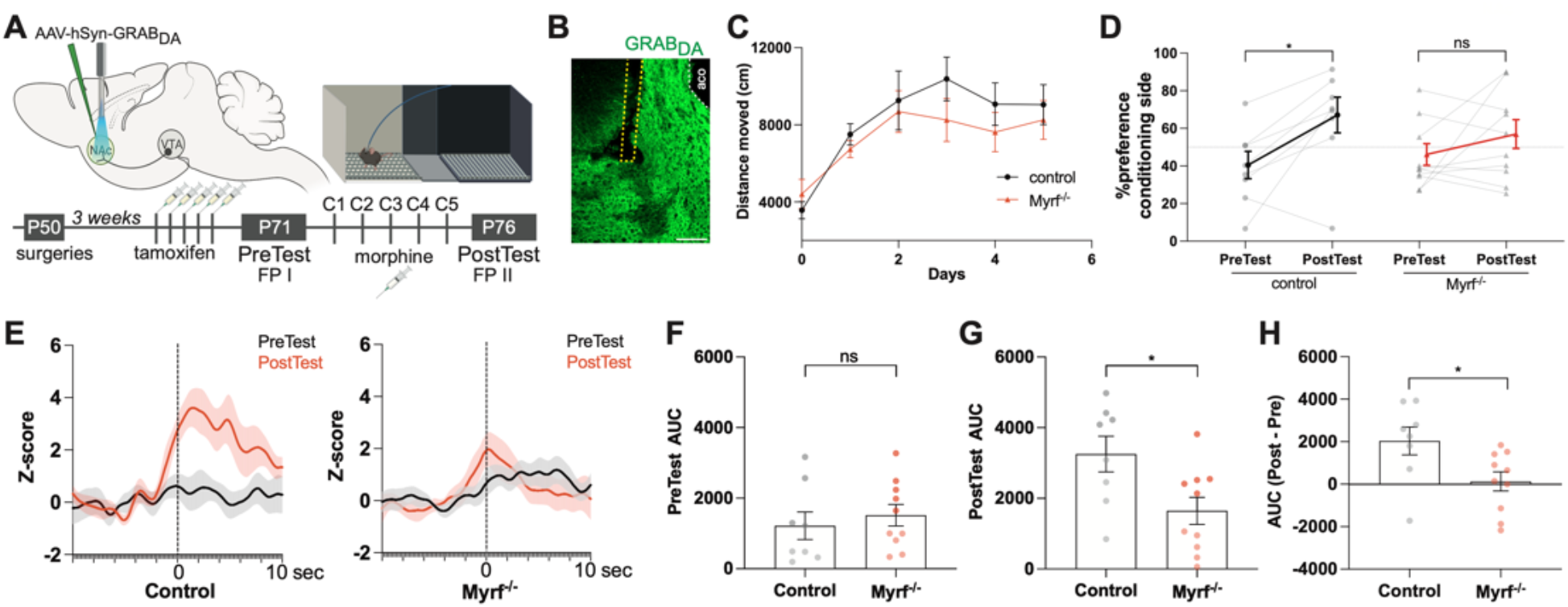
Morphine-induced oligodendrogenesis alters DA dynamics at NAc. **(A)** Schematic of experimental paradigm to determine the morphine conditioning effects in DA dynamics in *Myrf^-/-^* (*Myrf^-/-^; Pdgfra-Cre^ERT^)* and littermate controls; conditional knock-out performed with tamoxifen administration prior to CPP. DA dynamics measured with fiber photometry during PreTest and PostTest as mice freely explored CPP chambers. Mice groups injected with morphine (10mg/kg) during conditioning sessions. **(B)** Confocal microgram shows the fiber photometry cannula positioning in NAc outlined by yellow dashed lines, white dashed line marks anterior commissure (aco). GRAB_DA_ (green) expression in NAc below the cannula, scale bar= 100 μm. **(C)** Morphine increases locomotor sensitization in both control and *Myrf^-/-^* mice. **(D)** Control mice acquire a place preference for the morphine conditioning chamber, whereas *Myrf^-/-^* mice do not show a strong preference. Graph shows the percentage of time spent in conditioning chamber, comparing PreTest and PostTest. **(E)** Group average GRAB_DA_ responses upon first entry into the conditioning chamber. Control mice show increased DA release after morphine conditioning paradigm (left), in contrast, *Myrf^-/-^* mice do not exhibit a similar increase (right). **(F)** During PreTest, DA release in control and *Myrf^-/-^* mice are similar, while during **(G)** PostTest only control animals show increased DA release upon first entry into the conditioning chamber. **(H)** Control mice release more DA in PostTest compared to *Myrf^-/-^* mice (control, n=8 mice; *Myrf^-/-^*, n=10 mice). AUC (area under the curve), **D**, Wilcoxon matched-pairs test, **F, G, H,** unpaired two-tailed t-test, ns (not significant) p>0.5, *p<0.05. Data shown as mean, error bars indicate SEM.

## Discussion

Mounting evidence indicates that experience-dependent changes in myelin are necessary for proper neural circuit function and brain health. Here, we show that myelin plasticity occurs in the DA reward system and may become maladaptive in the context of opioid reward after repeated cycles of intoxication and withdrawal, thereby promoting the development of opioid use disorder.

Our findings suggest that only reward-related phasic (more than 10 Hz) DA neuron activity promotes an oligodendroglial response in VTA, not the cell-autonomous pacemaker tonic firing (0.2-10 Hz). Increasing phasic activity of DA neurons, either directly by optogenetic stimulation or indirectly by morphine administration, induced oligodendrogenesis specifically in VTA. Phasic firing is characterized by bursts of action potentials that result in synchronized activity across DA neurons that arises from shared inputs (*34*). Conceivably, phasic firing of DA neurons regulates myelin plasticity-mediated circuit adaptations to synchronize neuronal activity across the circuit and increase spatiotemporal precision - such as coordinated or enhanced DA release in NAc - to mediate reward-related behaviors. As a form of plasticity regulating VTA DA neuron function, this myelination is predicted to enhance synchronized DA release during cue exposure and facilitate the retrieval and expression of the reward association. Indeed, our fiber photometry recordings measuring real-time DA dynamics demonstrate that morphine-induced oligodendrogenesis is necessary for the precisely-timed DA release occurring when mice enter the previously paired morphine chamber, reflecting the learned association between the reward and the context (*49, 50*). Thus, blocking oligodendrogenesis before morphine conditioning attenuates the coordinated DA release in NAc and prevents associative learning in the conditioned place preference test. Thus, this previously unidentified form of plasticity in the VTA in response to morphine may be a critical factor in the powerful, maladaptive memories that contribute to the development of substance use disorders.

While blocking oligodendrogenesis prevented morphine CPP, it did not influence the development of locomotor sensitization indicating that this form of behavioral plasticity involves distinct mechanisms, perhaps occurring in the NAc (*51*). Indeed, we found that oligodendroglial cells proliferate in response to increased DA neuron activity specifically in the VTA, where the somas are located, but not in the major projection target NAc or along the DA fibers in MFB. This region-specific oligodendroglial change is also evident with the morphine-induced increases in new OPCs in VTA, concordant with the known effect of morphine to increase DA neuron activity (*51*). Even though the neurotransmitter DA or BDNF, a neurotrophin shown to mediate activity-regulated OPC proliferation, oligodendrogenesis and myelin plasticity in cortical projections (*29*), are released in both VTA and NAc upon neuronal activity (*52*), OPCs only respond in the VTA. This spatial specificity suggests region-specific mechanisms in VTA that may relate to particular populations of interneurons or other microenvironmental factors that future work will need to define. Regional heterogeneity amongst OPCs (*37, 38*) could also contribute to this selectivity.

While we demonstrate that DA neuronal activity promotes OPC proliferation and oligodendrogenesis, since oligodendroglial cells express k-opioid receptors (*40, 41*) morphine could exert direct effects on the oligodendroglial lineage in addition to the indirect effects exerted through opioid-induced increases in DA neuronal activity. While we did not detect a change in OPC proliferation in response to either DA or morphine alone *in vitro,* opioid pharmacology might still alter oligodendroglial dynamics *in vivo* where OPCs are presented with many additional regulatory factors and microenvironmental cues. In our experimental model, we chiefly assessed de novo oligodendrogenesis, and we cannot rule out the possibility that morphine might also cause direct differentiation of pre-existing OPCs. For example, methadone, an opioid that is commonly used as replacement therapy for opioid use disorder treatment, can act on oligodendroglial opioid receptors to increase myelination during development (*53*). In addition, dynorphin, an opioid neuropeptide related to stress that preferentially binds to k-opioid receptors, has been reported to promote oligodendrocyte differentiation (*54*). Future work delineating the effects of different opioid agents on oligodendroglial dynamics across the brain will provide a better understanding of how adaptive myelination is regulated normally and in individuals afflicted with substance use disorders.

Although a single dose of a drug is not sufficient to cause a substance use disorder, the resulting circuit alterations may lay the foundation for stronger and more persistent modifications that enable substance use disorder development. These persistent changes have been the focus of drug abuse research and are crucial for elucidating mechanisms that give rise to addictive behaviors. Here we show that morphine critically impacts myelin-forming oligodendroglial cells, driving maladaptive myelination of reward circuitry in the VTA which in turn regulates morphine-seeking behaviors. This study brings into focus another dimension of drug-evoked plasticity by defining new players in the neural circuit modifications that lead to addiction. Maladaptive myelination in the VTA could represent a key neural substrate of this pathological learning, suggesting myelination as a potential therapeutic target for opioid use disorder.

## Acknowledgments

We thank Anna Mathia Klawonn for helpful discussions and technical assistance. Funding: National Institute of Neurological Disorders and Stroke (R01NS092597 to MM), NIH Director’s Pioneer Award (DP1NS111132 to MM), National Institute for Drug Abuse (P50DA042012 to RCM and T32DA035165 and K99DA056573 to MBP), National Cancer Institute (P50CA165962, R01CA258384, U19CA264504 to MM), Robert J. Kleberg, Jr. and Helen C. Kleberg Foundation (to MM) Cancer Research UK (to MM), Maternal and Child Health Research Institute at Stanford University Postdoctoral Award (to BY), Dean’s Postdoctoral Fellowship at Stanford University (to BY)

## Author contributions

Conceptualization: BY, MM, Methodology: BY, MBP, RCM, MM, Investigation: BY, MBP, KM, IJC, KRT, LN, DC-E, Visualization: BY, Funding acquisition: MM, Project administration: BY, Supervision: MM, RCM, Writing – original draft: BY, MM, Writing – review & editing: BY, MBP, RCM, MM

## Data and materials availability

All data are available in the main text or the supplementary materials.

## Materials and Methods

### Animals

Both male and female mice were used equally in all experiments. Hemizygous Dat-Cre (The Jackson Laboratory, 006660) mice were bred with C57BL/6J (The Jackson Laboratory, 000664). Optogenetic experiments were performed on animals hemizygous for Dat-Cre. For conditional deletion of Myrf, hemizygous Pdgfra-CreER™ (The Jackson Laboratory, 018280) mice were bred with homozygous Myrf^lox/lox^ (The Jackson Laboratory, 010607) to generate hemizygous Pdgfra-CreER™ and heterozygous Myrf^lox/+^, which were backcrossed to homozygous Myrf^lox/lox^ to achieve hemizygous Pdgfra-CreER™ and homozygous Myrf^lox/lox^ (Myrf^lox/lox;^ Pdgfra-CreER™). Littermates that lack Pdgfra-CreER™ were used as control animals. To initiate Cre-dependent deletion of Myrf, animals were intraperitoneally injected with 100 mg/kg tamoxifen (Sigma-Aldrich) for 5 days at 7 weeks of age. For testing morphine effects on oligodendroglial cells, wild type C57BL/6J (The Jackson Laboratory, 000664) mice were used. All animals for housed in a 12-hour light/dark cycle with unrestricted access to food and water. All procedures were performed in accordance with guidelines set in place by the Stanford University Institutional Care and Use Committee (IACUC).

### Viral Vectors

For optogenetics experiments, Dat-Cre dependent expression of channelrhodopsin was achieved by injection of AAV-DJ-EF1a-DIO-hChR2(H134R)::eYFP (virus titer= 1.3×10^12^ gc/ml), and eYFP control by injection of AAV-DJ-EF1a-DIO-eYFP (virus titer= 1.6×10^12^ gc/ml). Both viral vectors were obtained from Stanford University Gene Vector and Virus Core. For fiber photometry experiments, GrabDA expression was achieved by injection of AAV9-hSyn-DA2m (DA4.4) (virus titer= 1.64×10^13^ gc/ml) obtained from WZ Biosciences (*55*).

### Surgical Procedures

Animals were anesthetized with 1-4% isoflurane and placed in a stereotaxic apparatus. For optogenetic stimulation experiments, 1 μl of AAV-ChR2::eYFP or AAV-eYFP viral vectors were unilaterally injected using Hamilton Neurosyringe and Stoelting stereotaxic injector over 5 minutes. Coordinates for viral injections and optic ferrule placements were measured from bregma. Viral vectors were injected into Ventral Tegmental Area (VTA) at coordinates, AP=-2.8mm, ML=-0.3mm, DV=-4.4mm, and an optic ferrule was placed at AP=-2.8mm, ML=-0.3mm, DV=-4.00mm. For Nucleus Accumbens (NAc) stimulations an optic ferrule was placed at AP= +1.25mm, ML=-0.75mm, DV=-4.00mm.

### Optogenetic Stimulations

Optogenetic stimulations were performed at least 3 weeks after the viral vector delivery, and 1 week after optic ferrule implantation. Freely moving animals were connected to a 473 nm diode-pumped solid-state laser system with a monofiber patch cord. Phasic dopaminergic neuron stimulation, for both neuronal cell bodies and axon terminals, was performed at 30 Hz, eight 5 ms pulses of 473 nm light delivery every 5 seconds at a light power output of 15mW (~ 475mW mm^-2^) from the tip of the optic fiber (200 μm core diameter, NA=0.22 - Doric lenses) (*36*). Tonic dopaminergic neuron stimulation was performed at 1 Hz, twenty-five 15 ms pulses of 473 nm light delivery every 1 minute (*35*). Acute optogenetic stimulation session lasted for 30 minutes for or phasic or tonic stimulations. Animals were injected intraperitoneally with 40 mg/kg EdU (5- ethynyl-2’-deoxyuridine; Invitrogen, E10187) before and after the session, and perfused 3 hours after the start of the stimulation. Chronic phasic optogenetic stimulations were performed for 10 minutes a day, for 7 days, with EdU injections for all stimulation sessions. Animals were perfused 4 weeks after the cessation of stimulations.

### Fiber Photometry

AAV9-hSyn-DA2m (DA4.4) was injected into NAc (AP= +1.25mm, ML=-0.75mm, DV=-4.40mm), and an optic fiber (400 μm core diameter, NA=0.48 - Doric Lenses) was placed just above the injection site (AP= +1.25mm, ML=-0.75mm, DV=-4.30mm). Mice were administered with tamoxifen 1 day to 4 weeks prior to starting CPP. After allowing 3 to 4 weeks for viral expression, GrabDA signal was recorded during the PreTest while the mice explored each chamber of the three-chamber CPP apparatus for 30 minutes. Mice then went through morphine conditioning for 5 days. The GrabDA signal was then recorded again during PostTest exploration for 30 minutes, in an identical manner to the PreTest.

Fiber photometry was performed as previously described (*56*). Data were acquired with Synapse software controlling an RZ5P lock-in amplifier (Tucker-Davis Technologies). GrabDA was excited by frequency-modulated 465 and 405 nm LEDs (Doric Lenses). Optical signals were band-pass-filtered with a fluorescence mini cube FMC4 (Doric Lenses) and signals were digitized at 6 kHz. Signal processing was performed with custom scripts in MATLAB (MathWorks). Briefly, signals were de-bleached by fitting with a mono-exponential decay function and the resulting fluorescence traces were Z-scored. Videos were analyzed for the first entry into the morphine conditioning chamber for each animal for the PreTest and PostTest. Peristimulus time histograms were constructed by taking the average of 20 sec epochs of fluorescence consisting of 10 sec before and 10 sec after the chamber entry, which is defined as time = 0. Before averaging, each epoch was offset such that the Z-score averaged from −10 to −1 sec equaled 0. Area under the curve was defined as the integral between 0 and +10 sec. The AUC difference score in was calculated as the (Posttest AUC - PreTest AUC) for each animal.

### Behavioral Analysis

#### Real-time place preference

Real-time place preference was performed in a two-chamber acrylic box (each chamber: 45 cm x 20 cm x 25 cm) without any additional contextual cues. Each mouse was connected to a 473 nm laser system with a monofiber patch cord, and gently placed into the middle section of the cage at the beginning of the test. One chamber of the cage was randomly assigned for optogenetic stimulation prior to the test. Optogenetic stimulation was turned on every time mouse entered into the pre-assigned chamber, and turned off if the mouse moved to the other chamber. Each session lasted for 20 minutes. Ethovision XT software (Noldus) was used to determine the time animals spent in each chamber after the test in an automated and condition-blinded manner.

#### Conditioned place preference

The experimental protocol was adapted from the previously described method (*42*). Conditioned place preference was performed in an acrylic rectangular three-chamber apparatus with two distinct chambers which are connected by a neutral corridor. The “grid chamber” included a 3-mm wide grids in the floor texture, and black stripes over white walls. The “spots chamber” included a floor with 10-mm diameter holes, and white walls. Both chambers were squares (18 cm x 18 cm x 18 cm), and connected by a 10-cm wide corridor (10 cm x 18 cm x 18 cm). The entrance to each grid or spots chamber could be closed off with a transparent acrylic divider. On day 1, mice were tested for baseline preference towards either chamber by placing them in the cage. After 30 minutes of free exploration between chambers with two different flooring, mice were returned to their home cage. On the morning of day 2 (conditioning day 1), all mice were administered with saline intraperitoneally and placed in the non-conditioning chamber with either flooring for 15 minutes, and returned to their home cage. The morning session was followed by conditioning afternoon session 4 hours after. In the afternoon session, a group of mice received saline and EdU (40 mg/kg) injections intraperitoneally, and placed in the conditioning chamber for 30 minutes. Another group of mice were administered with 10mg/kg morphine and EdU (40 mg/kg) intraperitoneally and placed in the conditioning chambers. These conditioning sessions were repeated on the following days 3, 4, 5, and 6 (total conditioning days 1-5). On day 7, mice were again allowed to freely move around the cage for 30 minutes. The time spent on each chamber with either type of grid was analyzed using Ethovision XT software (Noldus) in an automated and condition-blinded manner.

#### Novel-object recognition

The experimental protocol was adapted from the previously described method (*45*). Animals handled daily for the week leading up to the test for 5 minutes each day, and habituated to the experimental room. Mice were placed in the acrylic experimental cage (50 cm x 50 cm x 50 cm) to acclimatize for 10 minutes on the day before testing. On the day of the testing, mice were tested for anxiety by placing into the experimental cage for 10 minutes, and video recorded. If an animal spent less than 2 minutes in the center of the box (10 cm away from the walls), it was regarded as too anxious for the test and discarded from analysis. Mice were then placed in home cage for 5 minutes. For the training phase, mice were placed in the experimental chamber with two identical inanimate objects (approximately 5 cm in size). Each time the mouse was placed in the experimental chamber it was facing away from the objects towards the opaque walls. The mouse was allowed to explore these objects for 10 minutes, then was returned to its home cage. Both the experimental chamber and the objects were cleaned using 70% ethanol. For the novel object testing phase, which is performed 24 hours after the training phase, the mouse was returned to the cage to explore these objects for 10 minutes. One of these objects was returned to the experimental cage and a novel inanimate object of similar size was placed into the cage. The objects used as novel and familiar, and the position of the novel object were counterbalanced from trial to trial, animal to animal. All the objects used were initially tested to ensure there was no bias, or preference for the animals. All sessions were camera recorded and analyzed using Ethovision XT software (Noldus) in an automated and condition-blinded manner. Any exploratory head gesturing within 2 cm of the object, including sniffing and biting was considered as investigation, but climbing onto the object was not considered. Only animals that explored the objects for a minimum of 20 seconds were included in the analysis.

#### Sucrose preference

The experimental protocol was adapted from the previously described method (*46*). Mice were tested for sucrose preference 4 weeks after the induction of Myrf deletion by tamoxifen administration. On the first day, animals were habituated individually in experimental cages and drinking bottles with water. Next day, one water bottle, and one sucrose solution bottle (1% wt/vol) were provided for ad libitum access to mice. Water and sucrose solution were replenished every 24 hours, and consumption was determined per day over three days. Position of the sucrose and water bottles were swamped daily to prevent a bias towards bottle location. All measurements were corrected for spillage by subtracting the volume loss from a control bottle.

#### Three-chamber social interaction

Mice were tested for social preference 4 weeks after the induction of Myrf deletion by tamoxifen administration. Animals were habituated to the experimental room and the three-chamber experimental cage prior to test. Three-chamber acrylic experimental cage (each chamber: 45 cm x 20 cm x 25 cm) contained two metal grid pencil cups (10 cm in diameter) in each of the outer chambers. The chambers were divided with transparent acrylic walls with 15 cm wide entrance holes. One of the chambers contained a novel same-sex juvenile in the pencil cup, and the other chamber contained an inanimate object in the pencil cup. Test mouse was placed into the middle chamber and 1 minute after the chamber dividers were removed, and the mouse was allowed to explore each chamber freely for 20 minutes. The location of the novel mouse and the inanimate object was counterbalanced between sessions. The time spent on interacting with either pencil cup was analyzed using Ethovision XT software (Noldus) in an automated and condition-blinded manner.

### *In vitro* OPC Analysis

To test the direct effects of morphine and dopamine, in vitro OPC proliferation assay was used. C57BL/6 P4-5 mice pups were rapidly decapitated and brains were processed in Hibernate-A medium (Thermo Fisher Scientific, A12475-01). Resulting tissue was enzymatically disassociated in buffer containing HEPES-HBSS with DNase (Worthington Biochemical LS002007) and Liberase (Roche Applied Sciences 05401054001) at 37°C on a rotator. Tissue mixture then was triturated with 1000μl tip and passed through a 100 μm cell strainer. OPCs were isolated using the CD140 (Pdgfra) Microbead kit (MACS, Miltenyi Biotex 130-101-502) according to the manufacturer’s instructions. 30,000 cells were seeded per well, on laminin-coated (Thermo Fisher Scientific, 23017015) cover slips in 24-well plate. OPC media containing DMEM (Thermo Fisher Scientific, 11320082), glutamax (Invitrogen, 35050-061), sodium pyruvate (Invitrogen, 11360070), MEM non-essential amino acids (Thermo Fisher Scientific, 11140076), antibiotic-antimytotic (GIBCO), N21-MAX (R&D systems, AR012), trace elements B (Corning, 25-022-Cl), 5 mg/ml N-acetyl cysteine (Sigma-Aldrich, A9165), 10 ng/ml PDGFAA (Shenandoah Biosciences, 200-54), 10 ng/ml CNTF (PeproTech, 450-13), and 1 ng/ml NT-3 (PeproTech, 450-03) was used. For proliferation studies, after 3 days in proliferative media, cells were treated with various concentrations of morphine or dopamine for 24 hours. In the last 4 hours of this treatment, 10 μM EdU was added to the media to label dividing cells. Afterwards, cells were fixed in 4% PFA for 20mins, and incubated in HBSS until immunohistochemistry. All *in vitro* experiments were performed in triple wells (technical replicate) and independently replicated (biological replicates).

### Immunohistochemistry

All mice were anesthetized with intraperitoneal injections of 2.5% Avertin (tribromoethanol; Sigma-Aldrich, T48402), and transcardially perfused with 20 ml 0.1M phosphate buffer saline (PBS). Brains were postfixed in 4% paraformaldehyde (PFA) overnight at 4°C before cryoprotection in 30% sucrose solution for 48 hours. For sectioning, brains were embedded in optimum cutting temperature (OCT; Tissue-Tek) and sectioned coronally at 40 μm using a sliding microtome (Leica, HM450). For immunohistochemistry, brain sections were stained using the Click-iT EdU cell proliferation kit (Invitrogen, C10339) according to manufacturer’s protocol. Tissue sections were then stained with antibodies following an incubation in blocking solution (3% normal donkey serum, 0.3% Triton X-100 in tris buffer saline) at room temperature for 30 minutes. Goat anti-Pdgfra (1:500; R&D Systems, AF1062), rabbit anti-Olig2 (1:500; Abcam, 7349), rabbit anti-ASPA (1:250; EMD Millipore, ABN1698), rat anti-MBP (1:200; Abcam ab7349), rabbit anti-tyrosine hydroxylase (1:500; Millipore Sigma, AB152), mouse anti-tyrosine hydroxylase (1:250; Novus Biologicals, MAB7566), chicken anti-GFP (1:1000; Aves Labs, GFP-1020), chicken anti-mCherry (1:1000; ab205402), or rabbit anti-cfos (1:500; Santa Cruz Biotechnology, sc-52) were diluted in 1% blocking solution (1% normal donkey serum in 0.3% Triton X-100 in TBS) and incubated overnight at 4°C. All antibodies have been validated in the literature for use in mouse immunohistochemistry. To further validate the antibodies, we confirmed that each antibody stained in the expected cellular patterns and brain-wide distributions (e.g., nuclear Olig2 staining, cell membrane Pdgfra staining, tyrosine hydroxylase staining in midbrain dopaminergic neurons). The following day, brain sections were rinsed three times in 1x TBS and incubated in secondary antibody solution for 2 hours at the room temperature. All secondary antibodies were used at 1:500 concentration including Alexa 488 anti-rabbit (Jackson ImmunoResearch; 711-545-152), Alexa 488 anti-mouse (Jackson ImmunoResearch; 715-545-150), Alexa 488 anti-chicken (Jackson ImmunoResearch; 703-545-155), Alexa 594 anti-chicken (Jackson ImmunoResearch; 703-585-155), Alexa 647 donkey anti-goat (Jackson ImmunoResearch; 705-605-147), Alexa 647 anti-rat (Jackson ImmunoResearch; 712-605-150), Alexa 647 anti-rabbit (Jackson ImmunoResearch; 711-605-152), Alexa 647 donkey anti-goat (Jackson ImmunoResearch; 705-605-147). Sections were then rinsed three times in 1x TBS and mounted with ProLong Gold (Life Technologies, P36930).

### Fluorescence microscopy and quantification

All image analysis were performed by experimenters blinded to the experimental conditions or genotype. EdU stereology images were taken using a Zeiss AxioObserver upright fluorescence microscope with automated stage and tile-scanning capability (Stereo Investigator, Microbrightfield) with 20x objective. Imaging and cell counting were performed on every 6^th^ 40 μm slice, throughout the extend of dopaminergic neurons in the midbrain, and extend of NAc. Cells surrounding ferrule-induced tissue damage was not included. Regions with tyrosine hydroxylase labelling (or Dat-Cre>>GFP positive) was marked, and EdU cells within these regions were counted manually using Stereo Investigator software. Resulting numbers were reported as a function of the area (cells per mm^2^). Higher resolution imaging was conducted by acquiring z-stacks using a Zeiss LSM710 confocal microscope (Carl Zeiss). For MBP intensity analyses, z-stack images were taken of VTA using confocal microscopy with the same laser and fluorescence settings across animals and sections. Mean MBP intensity of pixels over VTA, which is labelled by Dat-Cre>>GFP, was quantified using Fiji and reported as a function of the area.

---

**Fig. S1.**
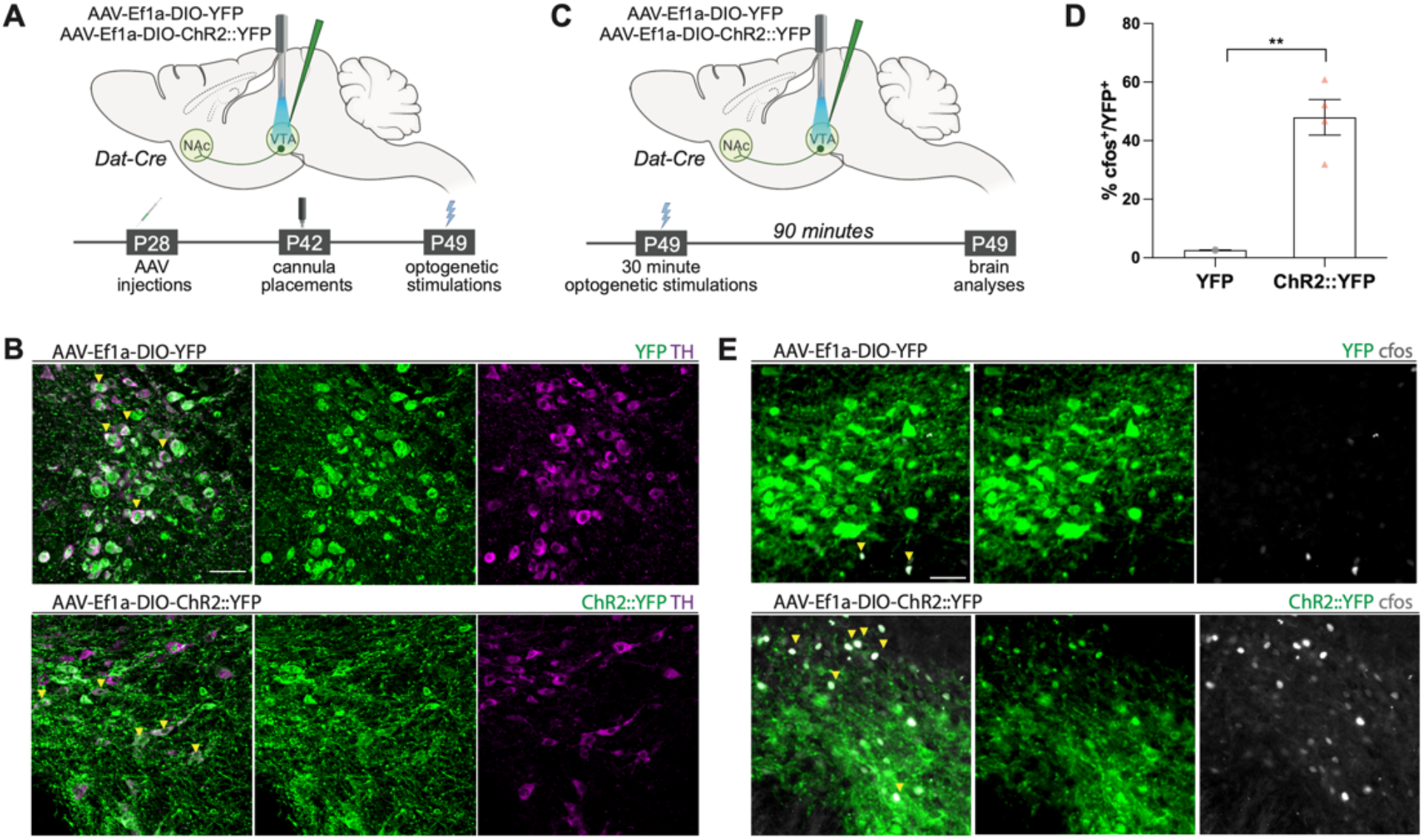
Optogenetic activation of DA neuron activity. **(A)** Schematic of experimental timeline for optogenetics experiments **(B)** Confocal micrographs show colocalization of YFP (green, panels above) or ChR2::YFP (green, panel below) with the DA neuron marker, TH (magenta) in VTA, scale bar= 50 μm **(C)** Schematic of experimental timeline for optogenetics stimulations. **(D)** 30 hz optogenetic stimulation of ChR2::YFP^+^ neurons increases cfos labeling in VTA within 90 minutes. **(E)** Confocal micrographs show colocalization of YFP (green, panels above) or ChR2::YFP (green, panel below) with the immediate early gene marker, cfos (gray) indicating neuronal activity in VTA. Unpaired two-tailed t-test, *p<0.05. Data shown as mean, error bars indicate SEM.

**Fig. S2.**
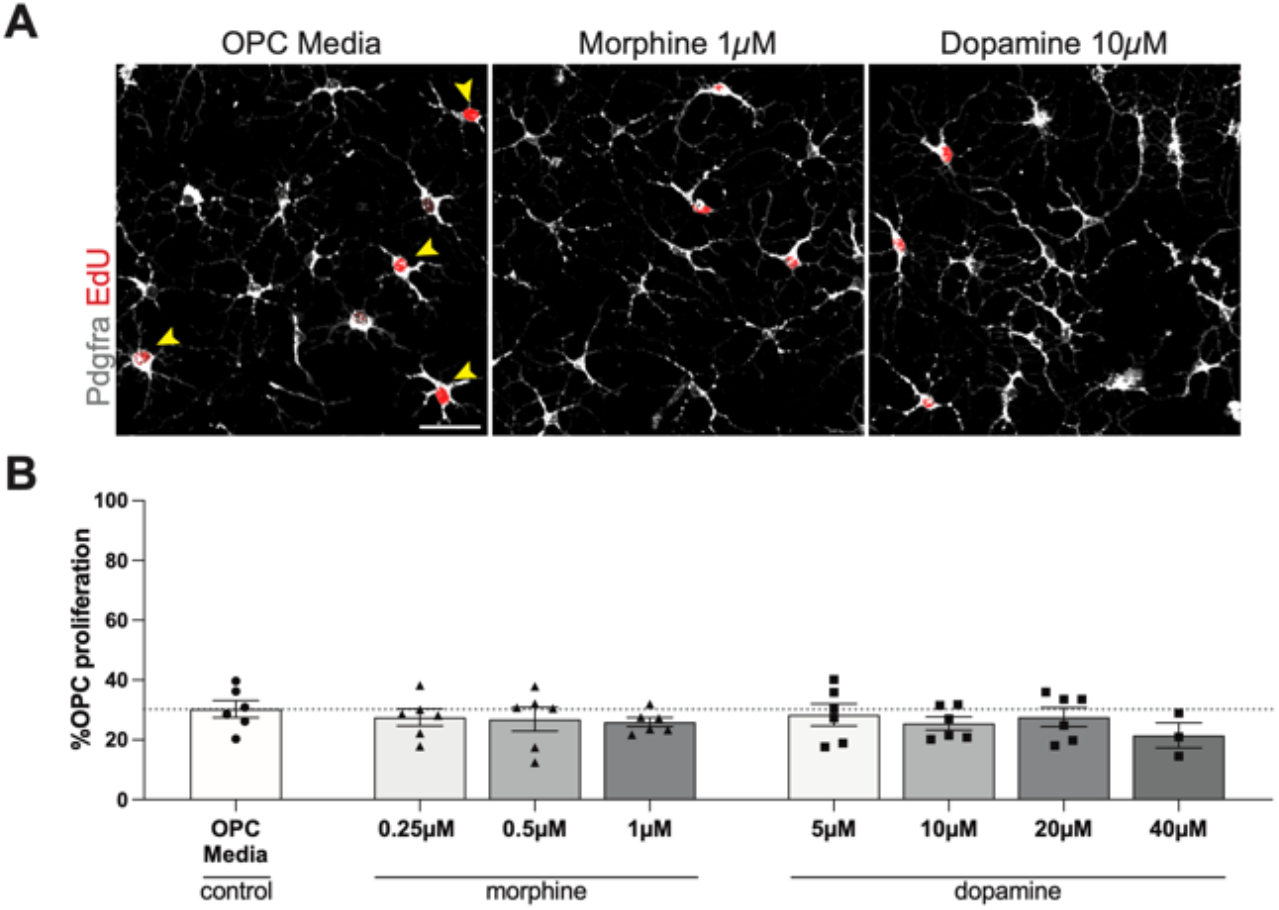
Morphine shows no direct effect on OPC proliferation *in vitro.* **(A)** Confocal micrographs show new OPCs (Pdgfra^+^ EdU^+^) in vitro when exposed to control OPC media, 1 μM morphine, or 10 μM dopamine. **(B)** Quantification of new OPCs (Pdgfra^+^ EdU^+^) after exposure to different concentrations of morphine and dopamine show similar OPC proliferation compared to OPC media control (each point indicates one technical replicate) Unpaired two-tailed t-test, *p<0.05. Data shown as mean, error bars indicate SEM.

**Fig. S3.**
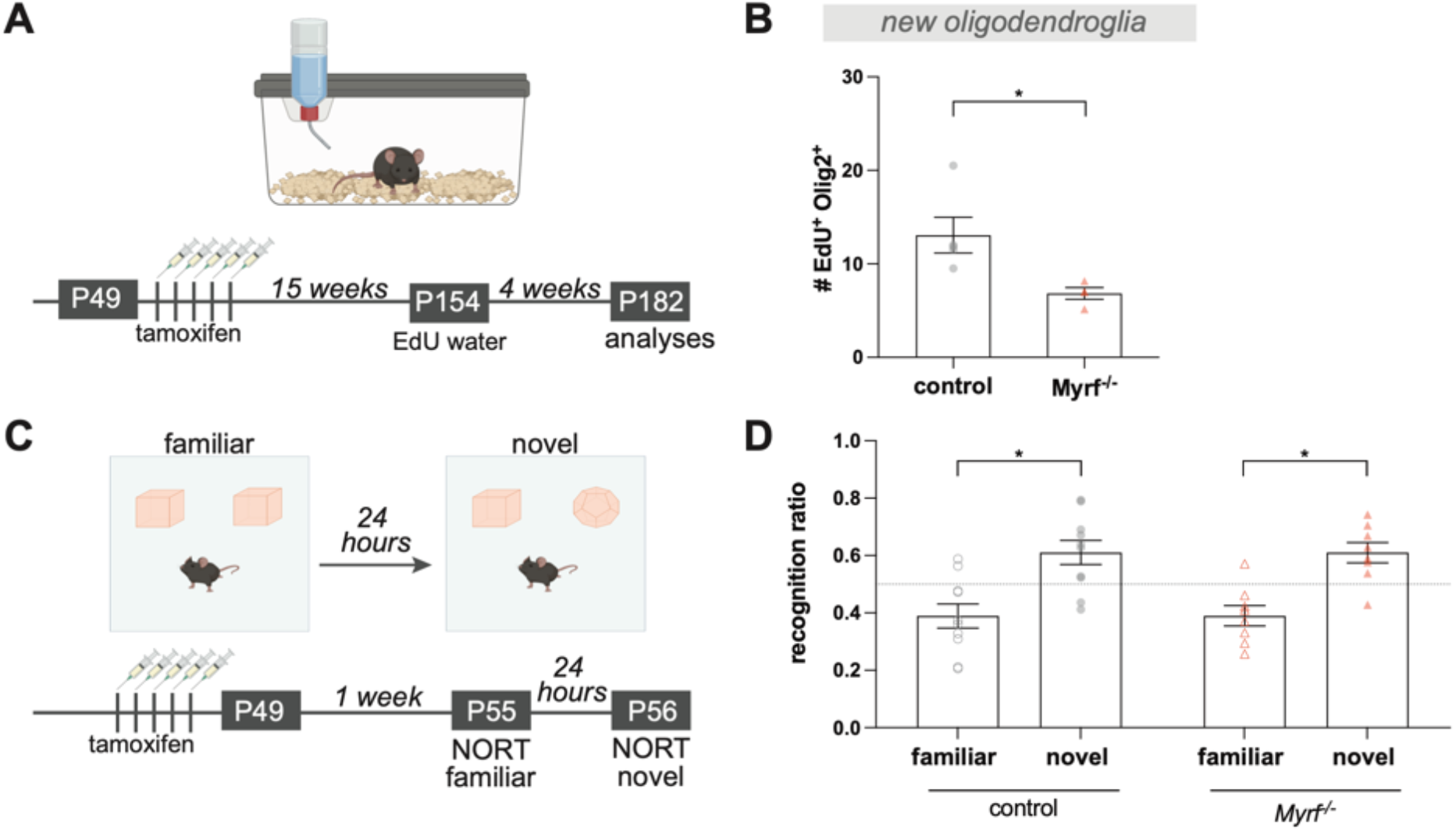
Myrf loss causes reduced oligodendrogenesis but not memory impairment. **(A)** Schematic of experimental paradigm for determining the effects of Myrf loss on oligodendrogenesis; 15 weeks after conditional knock-out of Myrf by tamoxifen administration in *Myrf^-/-^* (*Myrf^-/-^; Pdgfra-Cre^ERT^*) and littermate controls, EdU is added to drinking water to measure the baseline levels of oligodendrogenesis within 4 weeks. **(B)** *Myrf^-/-^* animals show reduced new oligodendroglia density in corpus callosum white matter compare to controls (control, n=5 mice; *Myrf^-/-^*, n=4 mice). **(C)** Schematic of experimental paradigm for novel object recognition test; mice are introduced to two identical objects and tested 24 hours after for memory by switch one of the objects with a novel object. 1 week after the end of tamoxifen administration *Myrf^-/-^* (*Myrf^-/-^; Pdgfra-Cre^ERT^*) and littermate controls are tested in novel object recognition test. This timeline is identical to CPP paradigms tested in Fig. 4A. **(D)** Both control and *Myrf^-/-^* and control animals spend more time with novel object compared to the familiar object (control, n=10 mice; *Myrf^-/-^*, n=8 mice). Paired two-tailed t-test for each genotype, *p<0.05. Data shown as mean, error bars indicate SEM.

## References and Notes

1. D. Saal, Y. Dong, A. Bonci, R. C. Malenka, Drugs of abuse and stress trigger a common synaptic adaptation in dopamine neurons. Neuron. 37, 577–582 (2003).

2. S. E. Hyman, R. C. Malenka, E. J. Nestler, Neural mechanisms of addiction: The role of reward-related learning and memory. Annu. Rev. Neurosci. 29, 565–598 (2006).

3. E. J. Nestler, C. Luscher, The Molecular Basis of Drug Addiction: Linking Epigenetic to Synaptic and Circuit Mechanisms (2019), doi:10.1016/j.neuron.2019.01.016.

4. C. Lüscher, R. C. Malenka, Review Drug-Evoked Synaptic Plasticity in Addiction: From Molecular Changes to Circuit Remodeling. Neuron. 69, 650–663 (2011).

5. J. A. Kauer, R. C. Malenka, Synaptic plasticity and addiction. Nat. Rev. Neurosci. 8 (2007), pp. 844–858.

6. M. D. Scofield, H. A. Boger, R. J. Smith, H. Li, P. G. Haydon, P. W. Kalivas, Gq-DREADD selectively initiates glial glutamate release and inhibits cue-induced cocaine seeking. Biol. Psychiatry. 78, 441–451 (2015).

7. G. M. Lewitus, S. C. Konefal, A. D. Greenhalgh, H. Pribiag, K. Augereau, D. Stellwagen, Microglial TNF-α Suppresses Cocaine-Induced Plasticity and Behavioral Sensitization. Neuron (2016), doi:10.1016/j.neuron.2016.03.030.

8. D. T. Kashima, B. A. Grueter, Toll-like receptor 4 deficiency alters nucleus accumbens synaptic physiology and drug reward behavior. Proc. Natl. Acad. Sci. U. S. A. 114, 8865–8870 (2017).

9. M. J. Lacagnina, A. M. Kopec, S. S. Cox, R. Hanamsagar, C. Wells, S. Slade, P. M. Grace, L. R. Watkins, E. D. Levin, S. D. Bilbo, Opioid Self-Administration is Attenuated by Early-Life Experience and Gene Therapy for Anti-Inflammatory IL-10 in the Nucleus Accumbens of Male Rats. Neuropsychopharmacology. 42, 2128–2140 (2017).

10. W. Xin, K. E. Schuebel, K. wing Jair, R. Cimbro, L. M. De Biase, D. Goldman, A. Bonci, Ventral midbrain astrocytes display unique physiological features and sensitivity to dopamine D2 receptor signaling. Neuropsychopharmacology. 44, 344–355 (2019).

11. B. Y. A. F. Huxley, A. D. R. Stampfli, Evidence for Saltatory Conduction in Peripheral Myelinated Nerve Fibres. J. Physiol. 108, 315–339 (1948).

12. R. S. Smith, Z. J. Koles, Myelinated nerve fibers: computed effect of myelin thickness on conduction velocity. Am. J. Physiol. 219, 1256–1258 (1970).

13. U. Fünfschilling, L. M. Supplie, D. Mahad, S. Boretius, A. S. Saab, J. Edgar, B. G. Brinkmann, C. M. Kassmann, I. D. Tzvetanova, W. Möbius, F. Diaz, D. Meijer, U. Suter, B. Hamprecht, M. W. Sereda, C. T. Moraes, J. Frahm, S. Goebbels, K.-A. Nave, Glycolytic oligodendrocytes maintain myelin and long-term axonal integrity. Nature. 485(2012), doi:10.1038/nature11007.

14. A. S. Saab, I. D. Tzvetavona, A. Trevisiol, S. Baltan, P. Dibaj, K. Kusch, W. Möbius, B. Goetze, H. M. Jahn, W. Huang, H. Steffens, E. D. Schomburg, A. Pérez-Samartín, F. Pérez-Cerdá, D. Bakhtiari, C. Matute, S. Löwel, C. Griesinger, J. Hirrlinger, F. Kirchhoff, K. A. Nave, Oligodendroglial NMDA Receptors Regulate Glucose Import and Axonal Energy Metabolism. Neuron. 91, 119–132 (2016).

15. S. Mensch, M. Baraban, R. Almeida, T. Czopka, J. Ausborn, A. El Manira, D. A. Lyons, Synaptic vesicle release regulates myelin sheath number of individual oligodendrocytes in vivo. Nat. Neurosci. 18, 628–630 (2015).

16. J. H. Hines, A. M. Ravanelli, R. Schwindt, E. K. Scott, B. Appel, Neuronal activity biases axon selection for myelination in vivo. Nat. Neurosci. 18, 683–689 (2015).

17. E. M. Gibson, D. Purger, C. W. Mount, A. K. Goldstein, G. L. Lin, L. S. Wood, I. Inema, S. E. Miller, G. Bieri, J. B. Zuchero, B. A. Barres, P. J. Woo, H. Vogel, M. Monje, Neuronal Activity Promotes Oligodendrogenesis and Adaptive Myelination in the Mammalian Brain. Science (80-.). 344, 1252304–1252304 (2014).

18. S. Mitew, I. Gobius, L. R. Fenlon, S. J. McDougall, D. Hawkes, Y. L. Xing, H. Bujalka, A. L. Gundlach, L. J. Richards, T. J. Kilpatrick, T. D. Merson, B. Emery, Pharmacogenetic stimulation of neuronal activity increases myelination in an axonspecific manner. Nat. Commun. 9, 306 (2018).

19. E. G. Hughes, J. L. Orthmann-Murphy, Myelin remodeling through experience-dependent oligodendrogenesis in the adult somatosensory cortex. Nat. Neurosci., doi:10.1038/s41593-018-0121-5.

20. M. Swire, Y. Kotelevtsev, D. J. Webb, D. A. Lyons, C. Ffrench-Constant, Endothelin signalling mediates experience-dependent myelination in the CNS. Elife. 8(2019), doi:10.7554/eLife.49493.

21. C. M. Bacmeister, H. J. Barr, C. R. McClain, M. A. Thornton, D. Nettles, C. G. Welle, E. G. Hughes, Motor learning promotes remyelination via new and surviving oligodendrocytes. Nat. Neurosci. 23, 819–831 (2020).

22. S. M. Yang, K. Michel, V. Jokhi, E. Nedivi, P. Arlotta, Neuron class-specific responses govern adaptive myelin remodeling in the neocortex. Science. 370(2020), doi:10.1126/science.abd2109.

23. S. Pajevic, P. J. Basser, R. D. Fields, Role of myelin plasticity in oscillations and synchrony of neuronal activity. Neuroscience. 276, 135–147 (2014).

24. R. Noori, D. Park, J. D. Griffiths, S. Bells, P. W. Frankland, D. Mabbott, J. Lefebvre, Activity-dependent myelination: A glial mechanism of oscillatory self-organization in large-scale brain networks. Proc. Natl. Acad. Sci. U. S. A. 117, 13227–13237 (2020).

25. S. Moore, M. Meschkat, T. Ruhwedel, A. Trevisiol, I. D. Tzvetanova, A. Battefeld, K. Kusch, M. H. P. Kole, N. Strenzke, W. Möbius, L. de Hoz, K. A. Nave, A role of oligodendrocytes in information processing. Nat. Commun. 11, 1–15 (2020).

26. M. Makinodan, K. M. Rosen, S. Ito, G. Corfas, A Critical Period for Social Experience-Dependent Oligodendrocyte Maturation and Myelination. Science (80-.). 337, 1357–1360 (2012).

27. J. Liu, K. Dietz, J. M. Deloyht, X. Pedre, D. Kelkar, J. Kaur, V. Vialou, M. K. Lobo, D. M. Dietz, E. J. Nestler, J. Dupree, P. Casaccia, Impaired adult myelination in the prefrontal cortex of socially isolated mice. 15 (2012), doi:10.1038/nn.3263.

28. I. A. McKenzie, D. Ohayon, H. Li, J. Paes de Faria, B. Emery, K. Tohyama, W. D. Richardson, Motor skill learning requires active central myelination. Science (80-.). 346, 318–322 (2014).

29. A. C. Geraghty, E. M. Gibson, R. A. Ghanem, J. J. Greene, A. Ocampo, A. K. Goldstein, L. Ni, T. Yang, R. M. Marton, S. P. Paşca, M. E. Greenberg, F. M. Longo, M. Monje, Loss of Adaptive Myelination Contributes to Methotrexate Chemotherapy-Related Cognitive Impairment. Neuron. 103, 250–265.e8 (2019).

30. P. E. Steadman, F. Xia, M. Ahmed, A. J. Mocle, A. R. A. Penning, A. C. Geraghty, H. W. Steenland, M. Monje, S. A. Josselyn, P. W. Frankland, Disruption of Oligodendrogenesis Impairs Memory Consolidation in Adult Mice. Neuron. 105, 150–164.e6 (2020).

31. S. Pan, S. R. Mayoral, H. S. Choi, J. R. Chan, M. A. Kheirbek, Preservation of a remote fear memory requires new myelin formation. Nat. Neurosci. 23(2020), doi:10.1038/s41593-019-0582-1.

32. J. K. Knowles, H. Xu, C. Soane, A. Batra, T. Saucedo, E. Frost, L. T. Tam, D. Fraga, L. Ni, K. Villar, S. Talmi, J. R. Huguenard, M. Monje, Maladaptive myelination promotes generalized epilepsy progression. Nat. Neurosci. 25, 596–606 (2022).

33. K. A. Nave, H. Ehrenreich, Myelination and oligodendrocyte functions in psychiatric diseases. JAMA Psychiatry. 71, 582–584 (2014).

34. S. B. Floresco, A. R. West, B. Ash, H. Moorel, A. A. Grace, Afferent modulation of dopamine neuron firing differentially regulates tonic and phasic dopamine transmission. Nat. Neurosci. 6, 968–973 (2003).

35. H.-C. Tsai, F. Zhang, A. Adamantidis, G. D. Stuber, A. Bonci, L. de Lecea, K. Deisseroth, Phasic Firing in Dopaminergic Neurons Is Sufficient for Behavioral Conditioning. Science (80-.). 324, 1080–1084 (2009).

36. L. A. Gunaydin, L. Grosenick, J. C. Finkelstein, I. V. Kauvar, L. E. Fenno, A. Adhikari, S. Lammel, J. J. Mirzabekov, R. D. Airan, K. A. Zalocusky, K. M. Tye, P. Anikeeva, R. C. Malenka, K. Deisseroth, Natural neural projection dynamics underlying social behavior. Cell. 157, 1535–1551 (2014).

37. S. O. Spitzer, S. Sitnikov, Y. Kamen, O. De Faria, Oligodendrocyte Progenitor Cells Become Regionally Diverse and Heterogeneous with Age. Neuron (2019), doi:10.1016/j.neuron.2018.12.020.

38. R. Marisca, T. Hoche, E. Agirre, L. J. Hoodless, W. Barkey, F. Auer, G. Castelo-Branco, T. Czopka, Functionally distinct subgroups of oligodendrocyte precursor cells integrate neural activity and execute myelin formation. Nat. Neurosci. 23, 363–374 (2020).

39. E. G. Hughes, J. L. Orthmann-Murphy, A. J. Langseth, D. E. Bergles, Myelin remodeling through experience-dependent oligodendrogenesis in the adult somatosensory cortex (2018), doi:10.1038/s41593-018-0121-5.

40. C. Du, Y. Duan, W. Wei, Y. Cai, H. Chai, J. Lv, X. Du, J. Zhu, X. Xie, Kappa opioid receptor activation alleviates experimental autoimmune encephalomyelitis and promotes oligodendrocyte-mediated remyelination. Nat. Commun. 7(2016), doi:10.1038/ncomms11120.

41. F. Mei, S. R. Mayoral, H. Nobuta, F. Wang, C. Desponts, D. S. Lorrain, L. Xiao, A. J. Green, D. Rowitch, J. Whistler, J. R. Chan, Identification of the kappa-opioid receptor as a therapeutic target for oligodendrocyte remyelination. J. Neurosci. 36, 7925–7935 (2016).

42. C. L. Cunningham, C. M. Gremel, P. A. Groblewski, Drug-induced conditioned place preference and aversion in mice. Nat. Protoc. 1, 1662–1670 (2006).

43. M. J. Thomas, C. Beurrier, A. Bonci, R. C. Malenka, Long-term depression in the nucleus accumbens: a neural correlate of behavioral sensitization to cocaine. Nat. Neurosci. 4, 1217–1223 (2001).

44. B. Emery, D. Agalliu, J. D. Cahoy, T. A. Watkins, J. C. Dugas, S. B. Mulinyawe, A. Ibrahim, K. L. Ligon, D. H. Rowitch, B. A. Barres, Myelin Gene Regulatory Factor Is a Critical Transcriptional Regulator Required for CNS Myelination. Cell. 138, 172–185 (2009).

45. M. Leger, A. Quiedeville, V. Bouet, B. Haelewyn, M. Boulouard, P. Schumann-Bard, T. Freret, Object recognition test in mice. Nat. Protoc. 8, 2531–2537 (2013).

46. M. Y. Liu, C. Y. Yin, L. J. Zhu, X. H. Zhu, C. Xu, C. X. Luo, H. Chen, D. Y. Zhu, Q. G. Zhou, Sucrose preference test for measurement of stress-induced anhedonia in mice. Nat. Protoc. 13, 1686–1698 (2018).

47. F. Sun, J. Zeng, M. Jing, J. Zhou, J. Feng, S. F. Owen, Y. Luo, F. Li, H. Wang, T. Yamaguchi, Z. Yong, Y. Gao, W. Peng, L. Wang, S. Zhang, J. Du, D. Lin, M. Xu, A. C. Kreitzer, G. Cui, Y. Li, A Genetically Encoded Fluorescent Sensor Enables Rapid and Specific Detection of Dopamine in Flies, Fish, and Mice. Cell. 174, 481–496.e19 (2018).

48. W. X. Pan, R. Schmidt, J. R. Wickens, B. I. Hyland, Dopamine cells respond to predicted events during classical conditioning: Evidence for eligibility traces in the reward-learning network. J. Neurosci. 25, 6235–6242 (2005).

49. M. W. Howe, P. L. Tierney, S. G. Sandberg, P. E. M. Phillips, A. M. Graybiel, Prolonged dopamine signalling in striatum signals proximity and value of distant rewards (2013), doi:10.1038/nature12475.

50. E. S. Calipari, R. C. Bagot, I. Purushothaman, T. J. Davidson, J. T. Yorgason, C. J. Peña, D. M. Walker, S. T. Pirpinias, K. G. Guise, C. Ramakrishnan, K. Deisseroth, E. J. Nestler, In vivo imaging identifies temporal signature of D1 and D2 medium spiny neurons in cocaine reward. Proc. Natl. Acad. Sci. U. S. A. 113, 2726–2731 (2016).

51. P. Leone, D. Pocock, R. A. Wise, Morphine-dopamine interaction: Ventral tegmental morphine increases nucleus accumbens dopamine release. Pharmacol. Biochem. Behav. 39, 469–472 (1991).

52. E. M. Nikulina, C. E. Johnston, J. Wang, J. P. Hammer, Neurotrophins in the ventral tegmental area: Role in social stress, mood disorders and drug abuse. Neuroscience. 282, 122–138 (2014).

53. A. A. Vestal-Laborde, A. C. Eschenroeder, J. W. Bigbee, S. E. Robinson, C. Sato-Bigbee, The Opioid System and Brain Development: Effects of Methadone on the Oligodendrocyte Lineage and the Early Stages of Myelination. Dev. Neurosci. 36, 409–421 (2014).

54. L. A. Osso, K. A. Rankin, J. R. Chan, Experience-dependent myelination following stress is mediated by the neuropeptide dynorphin. Neuron, 1–14 (2021).

55. F. Sun, J. Zhou, B. Dai, T. Qian, J. Zeng, X. Li, Y. Zhuo, Y. Zhang, Y. Wang, C. Qian, K. Tan, J. Feng, H. Dong, D. Lin, G. Cui, Y. Li, Next-generation GRAB sensors for monitoring dopaminergic activity in vivo. Nat. Methods. 17, 1156–1166 (2020).

56. B. D. Heifets, J. S. Salgado, M. D. Taylor, P. Hoerbelt, D. F. Cardozo Pinto, E. E. Steinberg, J. J. Walsh, J. Y. Sze, R. C. Malenka, Distinct neural mechanisms for the prosocial and rewarding properties of MDMA. Sci. Transl. Med. 11, eaaw6435 (2019)

